# Estimation of Pharmacokinetic Parameters of Continuous Intravenous Flucloxacillin infusion in the Outpatient Settings (PKFLUCLOX)

**DOI:** 10.1101/499848

**Authors:** N Lwin, Z Liu, M Loewenthal, P Dobson, JW Yoo, P Galettis, K Lai, J Martin

**Affiliations:** Department of Immunology and Infectious Diseases, John Hunter Hospital, Newcastle, NSW, Australia; Faculty of Medicine and Public Health, University of Newcastle, NSW, Australia; Hunter Medical Research Institute, Newcastle, NSW, Australia; Clinical Pharmacology, Department of Medicine, The Royal Children’s Hospital, Melbourne, Australia

**Keywords:** Flucloxacillin, pharmacokinetics, drug concentration, clearance, continuous infusion, Hospital in the home (HITH), Outpatient parenteral antibiotic therapy (OPAT)

## Abstract

Flucloxacillin, a beta-lactam antibiotic of the penicillin class, is considered first line therapy for methicillin sensitive *Staphylococcus aureus* (MSSA) in Australia. (1). At our tertiary referral hospital in the home (HITH) program, it has been prescribed in a standard dosage of 8 grams per day by continuous infusion for more than 20 years. The aim of this observational study was to characterize the pharmacokinetic profile of flucloxacillin in patients who receive continuous infusion in the HITH setting, and to undertake population pharmacokinetic analysis performed with NONMEM software by comparing various structural models. This study utilised flucloxacillin concentrations from 44 separate specimens obtained from 23 patients. Twenty-five of these were collected immediately after elastomeric device removal, representing steady-state concentrations, and the remaining 19 were each collected at least 45 minutes after device removal to determine clearance of the drug. Plasma concentrations ranged from 13 to 194 mg/L with median steady-state concentration of 51.5mg/L and inter-quartile range of 24.6 mg/L. The time-course of flucloxacillin was best described by a 1-compartment model. The best three covariates, CrCL (ΔOFV= -11.7), eGFR (ΔOFV= -5.9) and serum albumin (ΔOFV= -5.8) were found to be equivalent in terms of decreasing the OFV. CrCL was superior in explaining inter individual variability. The best model for flucloxacillin clearance was a one compartment model with CrCL as the sole covariate. The estimated population parameters were 9.5 L for volume of distribution and 8.1 L/h for flucloxacillin clearance.

## Introduction

Administration of intravenous antibiotics at home in various Hospital in the Home (HITH) programs, also known as OPAT, Outpatient Parenteral Antibiotic Therapy, is now standard care once a patient has become medically stable. Despite this, the pharmacokinetics (PK) and the relationship of PK to pharmacodynamics (PD) of flucloxacillin has been minimally studied in this setting (2). Published studies have focused on comparing intermittent to continuous dosing in healthy volunteers and in critical illness. (3,4) These studies demonstrate marked variability in individual patient pharmacokinetic parameters and provide a case for therapeutic drug monitoring (TDM) in the setting of critical illness. (4) Therefore, estimation of the pharmacokinetic parameters associated with this form of dosing is needed. This information will then provide a rational basis for dosing and determine the requirement for therapeutic drug monitoring in thousands of patients every year.

## Background

Flucloxacillin is considered first line therapy for serious methicillin sensitive *Staphylococcus aureus* in Australia (1). Being a penicillin derivative, it exhibits “time-dependent” bacterial killing and has minimal “post-antibiotic effect” (5). This means that a concentration of flucloxacillin well above the minimum inhibitory concentration (MIC) for the relevant organism will be more effective than a concentration that intermittently falls below the MIC. Intermittent high concentrations will also not compensate for this. Stable concentrations above MIC are best achieved by continuous infusion. *Staphylococcus aureus* isolates with an MIC >2 mg/L are considered resistant. (6)

Our institution has operated a HITH program since 1995 and during that time has administered more than 45,500 days of intravenous flucloxacillin therapy via continuous infusion. The vast majority have received flat dose of 8gram daily. In practice, dose adjustments are only made for children and patients with severe renal impairment (e.g. glomerular filtrate rate (GFR) estimated to be less than 30mL/min.

## Aims

We aimed to describe the PK of continuous flucoxacillin infusion in patients with Staphylococcal infection. Using this PK data, we aimed to develop the simplest possible model of plasma flucloxacillin concentration during HITH that would both inform dosage and quantify inter-individual variation to determine the circumstances TDM is most appropriate. Such a model would potentially take into account the readily measurable quantities of age, sex, weight, height, renal function, and plasma albumin.

## Methods

Patients who were enrolled in our facility’s ‘Hospital-in-the-Home’ (HITH) service (the “Out & About” program) and who are receiving flucloxacillin were offered enrolment in our study provided that they were at least 18 years of age and not pregnant or breastfeeding.

Informed consent was obtained by one of the investigators. The study was approved by Hunter New England Human Research Ethics Committee. (HNEHREC Reference No: 17/05/17/4.01)

Flucloxacillin was administered by continuous infusion from an elastomeric device (“Mobifuser”, Slade Pharmacy, Mt. Kuring-gai, Australia) via a peripherally-inserted central catheter (PICC).

In some cases, patients were recruited a few days before the Out and About admission when the investigators identified them as suitable candidates. One patient was transferred to a HITH program in another district but was included in the study whilst awaiting transfer as they had already commenced their continuous flucloxacillin infusion in our facility. Patients had weekly assessment by a medical officer along with regular weekly haematological and biochemical parameters monitoring. Our HITH service has a nurse and a physician on call at all times to deal with any immediate problems.

### Specimen collection for flucloxacillin concentration measurements

Whilst on the Out & About program patient’s devices were changed daily. The first blood sample was taken at the time of device disconnection. After at least 45 minutes a second blood sample was drawn, and the exact time recorded. All measurements reported in this study used blood collected by peripheral venepuncture.

### Sample handling and preparation

At each time point, 5 ml of blood was collected into an EDTA blood tube. The blood was kept on crushed ice in an insulated container and transferred to the laboratory at Hunter Medical Research Institute (HMRI) by one of the investigators. The samples were centrifuged at 2000 x g for 10 minutes at 4°C. Following this plasma was aspirated from the sample using a sterile pipette. 1ml aliquots in 1ml volumes were placed into appropriately labelled vials. The aliquoted plasma was then stored in a - 80°C freezer. Bloods were processed within 1 hour of collection. Flucloxacillin was measured using liquid chromatography – mass spectrometry as described in Zhang et al (7).

### Data Collection

The following personal demographic, medical, and physical data were collected after enrolment or on the first study visit and recorded on the Case Record Form (CRF): height, weight, body mass index, date of birth, sex, diagnosis, dosage and duration of flucloxacillin, plasma creatinine and creatinine clearance using Cockcroft-Gault formula, and plasma albumin.

### PK analysis

Flucloxacillin was measured using liquid chromatography – mass spectrometry as described in Zhang et al (7). Briefly, 50 μL of sample plus 50 μL of Internal Standard was extracted with 400 μL acetonitrile. The sample was vortexed and the centrifuged at 15,000 xg for 5min. An aliquot was then transferred to a vial and injected into a Shimadzu 8060 LCMS system. The samples were analysed on a Kinetex C18 2.1 x 50mm (1.7μm) column with an initial mobile phase of 0.1% formic acid: methanol (70:30) for 0.5min followed by a linear gradient for 2 min to 0.1% formic acid: methanol (10:90) and held for 0.5min. The column was then re-equilibrated at the initial mobile phase for 0.5min. The sodium adduct of flucloxacillin was detected using positive electrospray at the following SRM’s 476-182, 476-182. Flucloxacillin eluted at 2.2 min and the calibration curve ranged from 0.1 – 100 mg/L.

### Population PK model development

NONMEM Version 7.2.0 (8) in combination with the gfortran compiler, PsN (Perl-speaks-NONMEM) version 4.7.0 (9,10), PLT tool (11), and R Version i386 3.3.1 (12) were used for model development and graphical illustration. Parameter estimation was conducted with the first-order conditional estimation with interaction (FOCE+I).

The published structural pharmacokinetic models for flucloxacillin include 1, 2, and 3 compartment models (13,14,15). We considered all these models in our study. Different error models including additive, proportional and combined error models were tested to describe the residual unexplained variability (RUV) of flucloxacillin. Inter-individual variability (IIV) as an exponential relationship was added and tested step-wise for each pharmacokinetic parameter. The off-diagonal elements, representing the covariance of IIVs were tested. Covariates with mechanistic plausibility were tested in this study including gender, age, weight, height, body mass index (BMI), estimated glomerular filtration rate (eGFR), creatinine clearance using Cockcroft-Gault formula (CrCL), and serum albumin. For selection of covariates, a stepwise forward selection followed by backward elimination procedure was applied. The likelihood ratio test was used to discriminate between alternative models. For the forward selection (covariate addition), a decrease in the objective function value (OFV) of 3.84 units (X^2^, p<0.05) in, was considered statistically significant for one parameter in a nested model. For the backward elimination (covariate subtraction), difference of OFV greater than 10.83 (X^2^, p<0.001) for deletion from the model was considered statistically significant. Reduction of IIV was also used to assess the influence of covariates. IIV shrinkage was taken into account for precision and bias metrics.

Conditional weighted residuals (CWRES) versus time (16), prediction-corrected and variance-corrected visual predictive checks (VPC) (17) were used as diagnostics. 95% bootstrap confidence intervals for the final population model parameters were conducted using PLT tools with subject replacement of 500 runs. (18)

## Results

### Synopsis

Median plasma flucloxacillin concentration at steady-state was 52 mg/L with the lowest concentration being 13 mg/L, and the patient with the highest CrCL of 253 mL/min. The best model for flucloxacillin clearance was a one compartment model with CrCL as the sole covariate with an additive random effect for inter-individual variability. Model diagnostics suggest that this model is both robust and reliable.

The estimated population parameters (table 3) were 9.5 L for volume of distribution and 8.1 L/h (135mL/min) for flucloxacillin clearance corresponding to a half-life of 49 minutes.

### Patient Characteristics

Twenty-three patients (18 males, 5 females) with a median age of 67 were included in this study and 2 of those participated twice (Table 1, Supplementary Table 4). All patients were being treated for serious staphylococcal infection. MSSA was identified in 18 patients (78 %) and the other five patients isolated coagulase negative staphylococci (Two *S. epidemidis*, Two *S.lugdunesis*, One *S. capitis*) which were sensitive to methicillin/flucloxacillin. Diagnoses included vertebral osteomyelitis (*n* = 2), bone and joint arthroplasty related infection (*n*=7), other bone and joint infection except vertebral osteomyelitis and arthroplasty (*n*=9), and bacteraemia with no deep source (*n*=5) (Table 2). Organisms were isolated in all 23 patients. No significant haematological and biochemical complications were noted. In particular, no patients developed flucloxacillin-associated neutropenia or hepatitis.

**Table 1.**
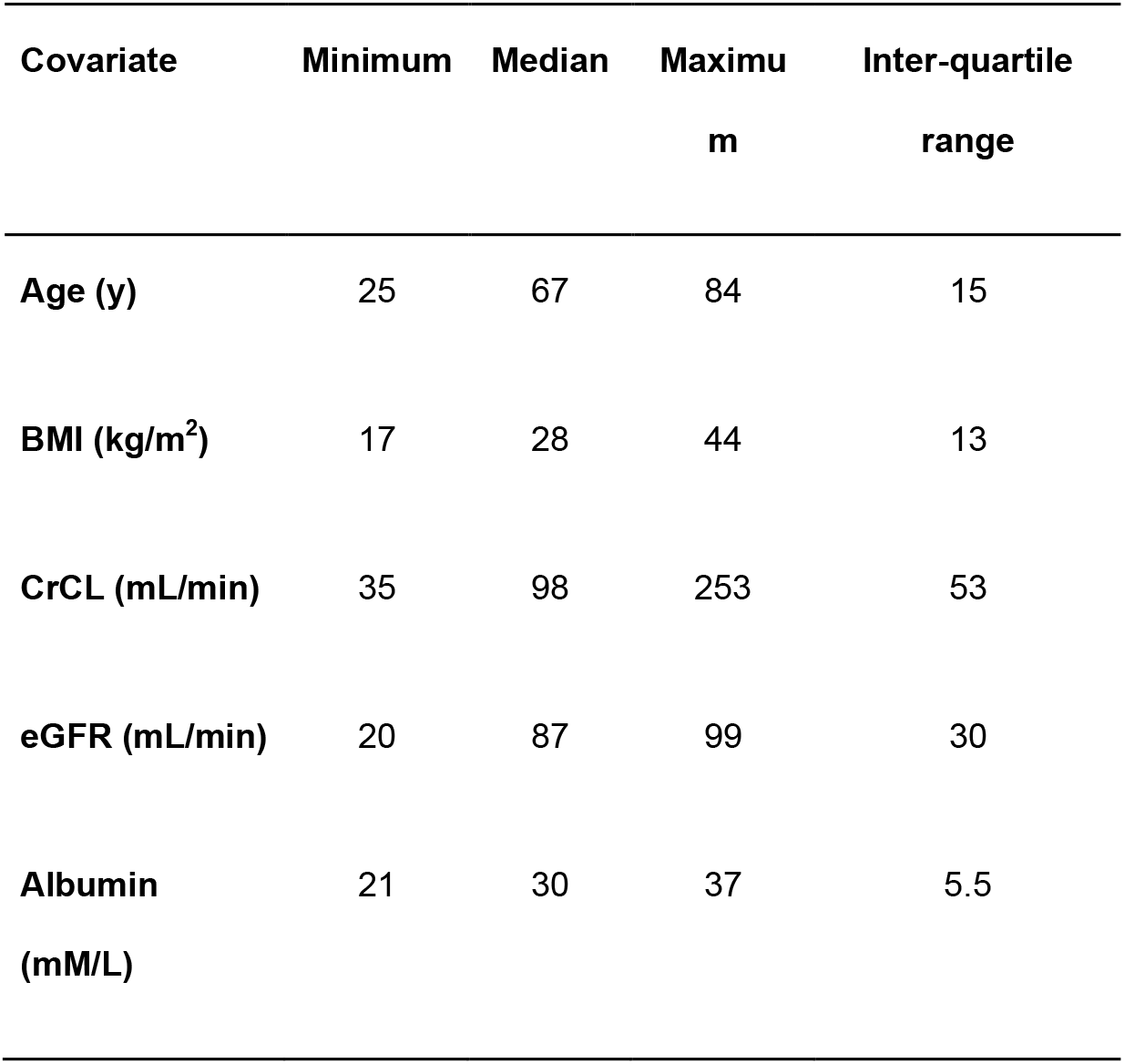
Patient characteristics. Body mass index (BMI), calculated creatinine clearance (CrCL), estimated glomerular filtration rate (eGFR).

**Table 2.** Diagnosis and reason for receiving flucloxacillin.

| Diagnosis | Number |
| --- | --- |
| Infected arthroplasty | 7 |
| Vertebral osteomyelitis | 2 |
| Other bone or joint infection | 9 |
| MSSA bacteraemia alone | 5 |

### Flucloxacillin Dosage

One patient received 6g/day of flucloxacillin. All other patients received 8g/day of flucloxacillin.

### Flucloxacillin concentrations

This study utilises flucloxacillin concentrations from 44 separate specimens. 25 of these were steady-state concentrations collected immediately after device removal. The remaining 19 were each collected at least 45 minutes after device removal. The lowest steady-state concentration in any patient was 13 mg/L. This was in the patient with the highest CrCL of 253 mL/min. The median steady-state concentration was 52 mg/L with a maximum of 194 mg/L and an inter-quartile range of 24.6 mg/L (Figure 1).

**Figure 1.**
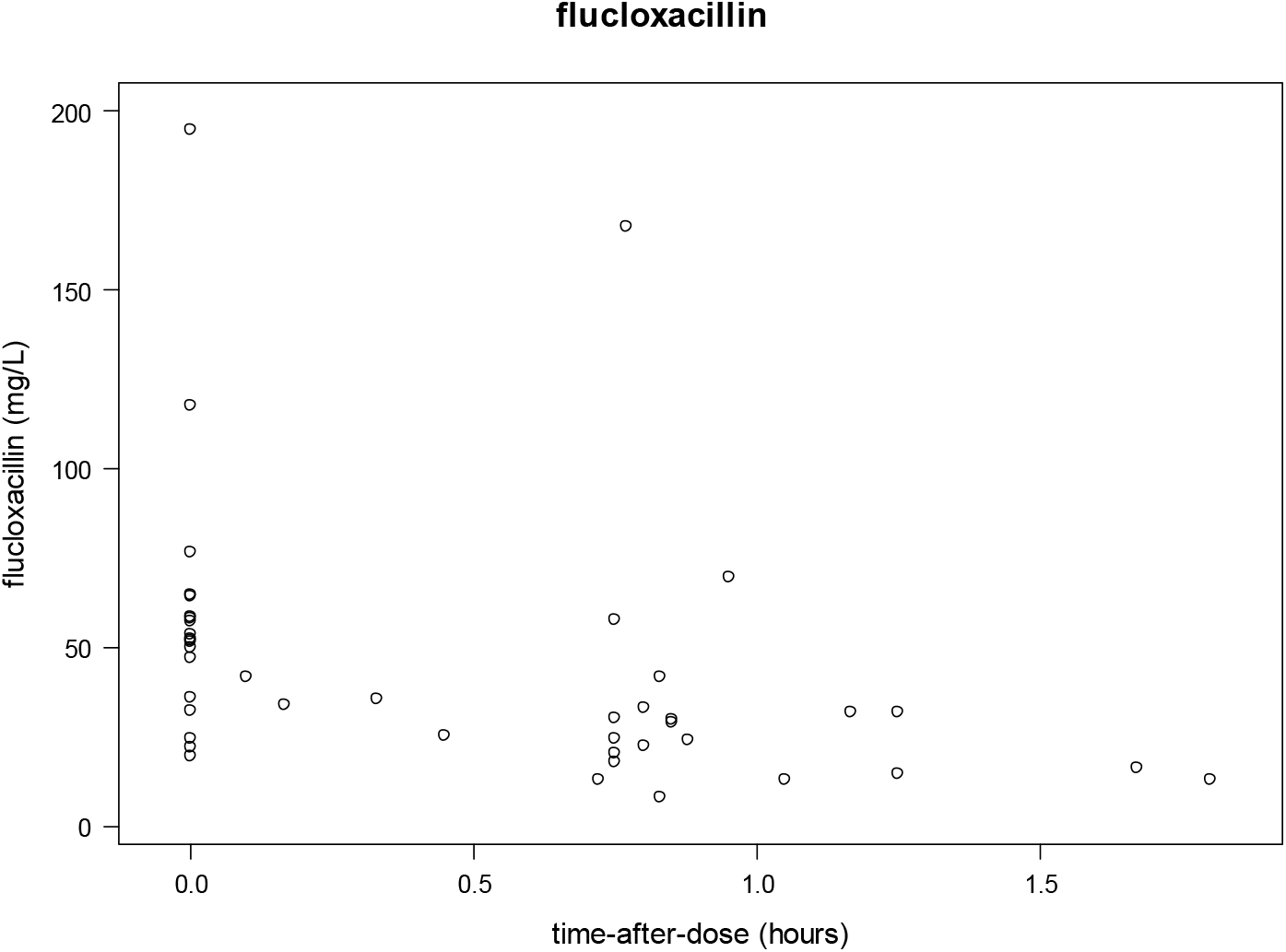
Observed concentrations (mg/L) versus time (hour) after cessation of infusion for flucloxacillin.

### Model development

The time-course of flucloxacillin was best described by a 1-compartment model. An additive residual error model was most appropriate to describe the RUV of flucloxacillin. The best final model included IIV on clearance. The best three covariates, CrCL (ΔOFV= -11.7), eGFR (ΔOFV= -5.9) and serum albumin (ΔOFV= - 5.8) were found to be equivalent in terms of decreasing the OFV. However, CRCL (ΔIIV= -11.6%) was superior to eGFR (ΔIIV= -6.7%) and serum albumin (ΔIIV= - 2.3%)in explaining the IIV. Therefore, in the final model CrCL was used as the covariate of clearance.

Conditional weighted residuals versus time after dose for the final model are presented in Figure 2. The VPCs are in Figure 3. These figures supported the model building process.

**Fig. 2.**
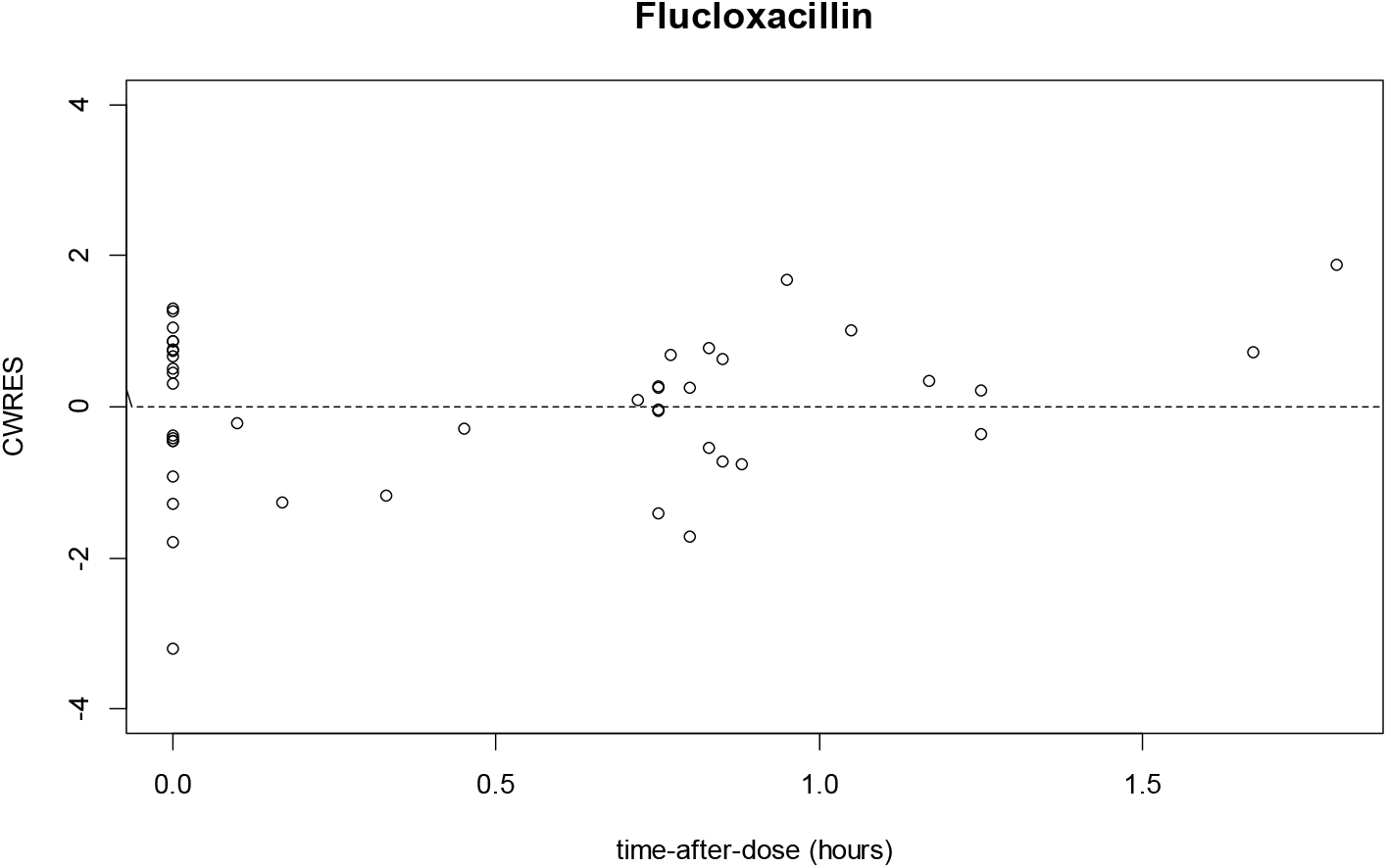
Conditional weighted residuals (CWRES) versus time after dose (hour) for flucloxacillin.

**Figure 3.**
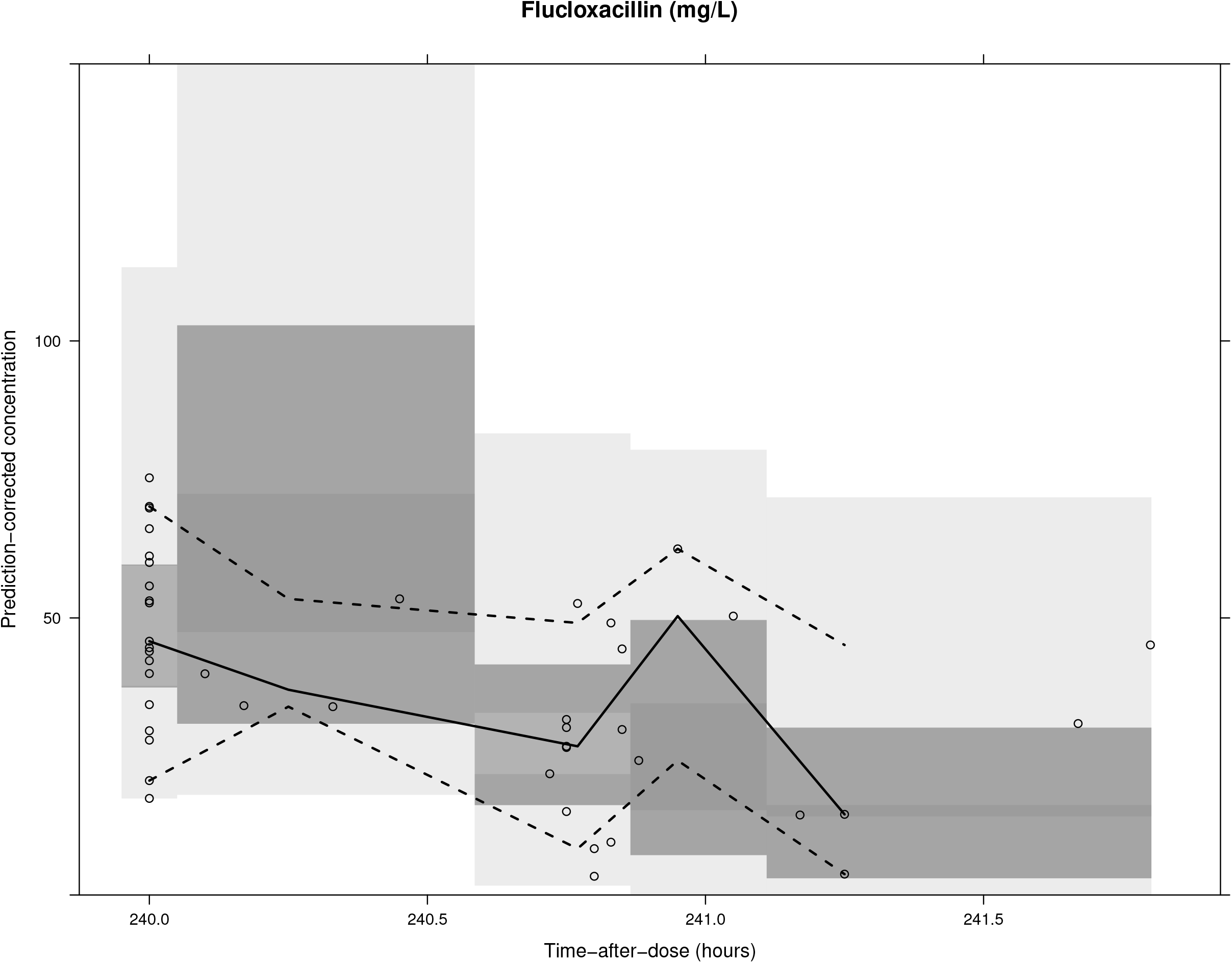
Prediction-corrected and variance-corrected VPCs for flucloxacillin. The raw data are represented as *black circles* and *lines*. Grey shaded areas are the 90% confidence interval for the 5^th^, 50^th^ and 95^th^ percentile of the simulated data. Five bins are used.

The final model parameters were also evaluated with a non-parametric bootstrap, which produced similar parameter values to those in the final model (Table 3), suggesting that the estimated parameter values in the final model are robust and reliable.

**Table 3.**
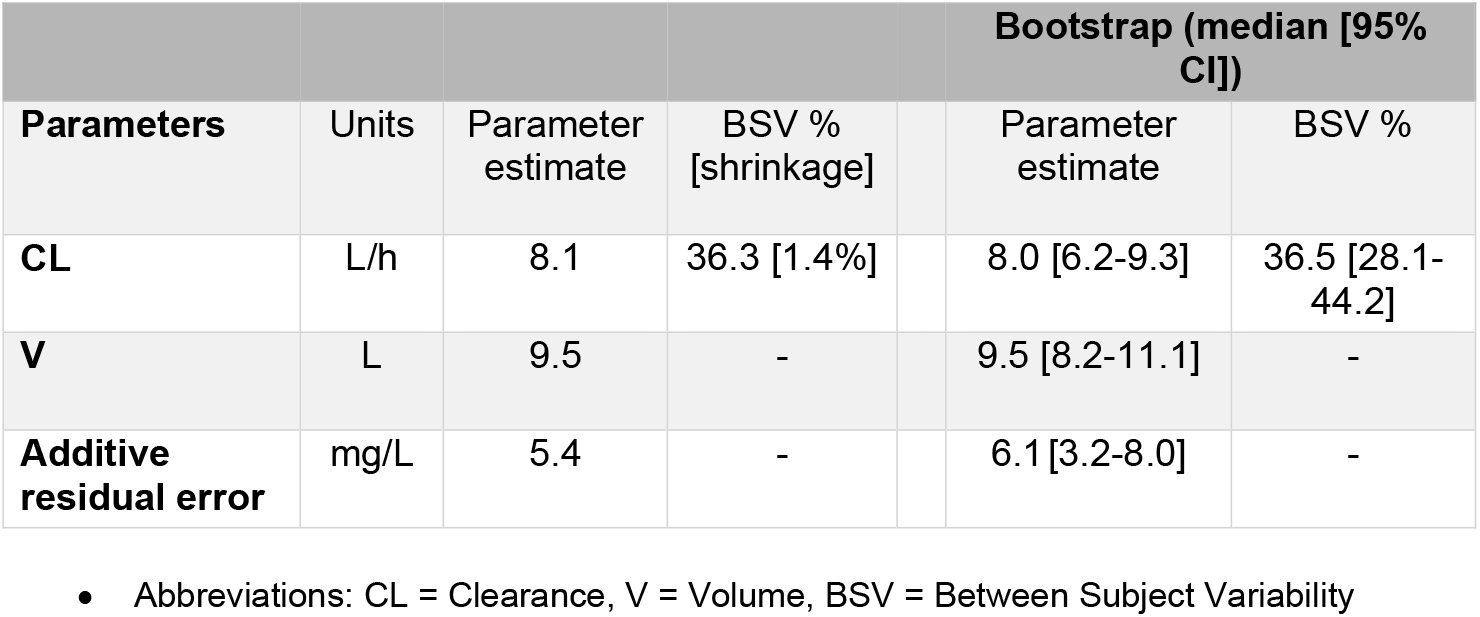
Estimated parameter values and bootstrap results for the final flucloxacillin model.

### Discussion

Our study used population pharmacokinetic modelling to estimate steady state flucloxacillin concentration in patients receiving this antibiotic by continuous infusion in the home setting. Achieving concentrations of β-lactam agents constantly at least 4 – 5 times the MIC produces the maximum bactericidal effect in the previous study (19).

Landersdorfer et al used a total daily dose of 6g in healthy volunteers and found that prolonged infusion and continuous infusion both had a 90% probability of producing concentrations above 1mg/L for more than 50% of the time whereas intermittent infusion at the same total daily dose could only do this for concentrations of 0.25 to 0.375 mg/L or less. (20). For deep-seated infection such as osteoarticular infection or endocarditis, 12 grams/day intermittent dosing is recommended (21). We estimated that the population half-life flucloxacillin to be 49 minutes, allowing concentrations to fall by 30-fold during a four-hour dosing interval. On this basis a continuous infusion of 8g daily should produce more time above the MIC than 12g daily by intermittent dosing. Our study revealed that a smaller dose of 8 grams daily by continuous infusion maintained the desired drug level (median=52 mg/L, minimum 13 mg/L), well above the MIC at all times. It follows that this dosing should be at least as effective as 12g daily intermittent dosing.

There is minimal information regarding antibiotic dosing in obese patients, and some suggest that beta-lactam antibiotics should be given at the upper end of the recommended dosage. (22) In our study, however, drug concentration did not appear to be independently associated with patients’ body mass indexes. Creatinine clearance is found to be superior to other covariates such as age, BMI, eGFR or serum albumin in predicting the flucloxacillin concentration.

Lastly, our study provides population estimates for essential pharmacokinetic parameters of flucloxacillin concentration at steady-state during continuous infusion via elastomeric device in the home setting. This finding helps inform future studies on topics such as the role of therapeutic drug monitoring, clinical outcomes and adverse reactions that may be related to drug concentration in the HITH setting. Imani et al. reported that patients being treated with intravenous flucloxacillin (given continuously or intermittently) who developed neurotoxicity were found to have minimum concentration of flucloxacillin with ≥ 125.1mg/L). (23) Our results suggest that patients receiving flucloxacillin via continuous intravenous infusion on a HITH program whose CrCL is between 39 and 200 mL/min can safely be given a standard dose of 8g over 24 hours.

#### Limitations

This is a single-centre study in which the number of study participants was small and only a limited number of covariates could be evaluated for their effect on population pharmacokinetic parameter estimates. Second, our study is designed to examine the pharmacokinetic properties of flucloxacillin and did not examine clinical outcomes.

### Conclusions

Continuous administration of flucloxacillin at dose of 8g over 24 hours using elastomeric devices on a home IV therapy program achieved plasma concentrations of well above five times the MIC for Staphylococci in all cases. CrCL calculated by the Cockgroft-Gault formula was the best predictor of flucloxacillin clearance. Although our study was small, it included patients with CrCL above 250 mL/min.

Median volume of distribution was estimated to be 9.5L and flucloxacillin clearance was estimated at 8.1L/hr (Table 3) based on population pharmacokinetic modelling which is consistent with studies using intermittent dosing.

Clinicians should be comfortable using this method of drug delivery. Consideration can be given to increasing the flucloxacillin dose only in patients with extremely high CrCL>250ml/min).

---

**Supplementary Table 4:**
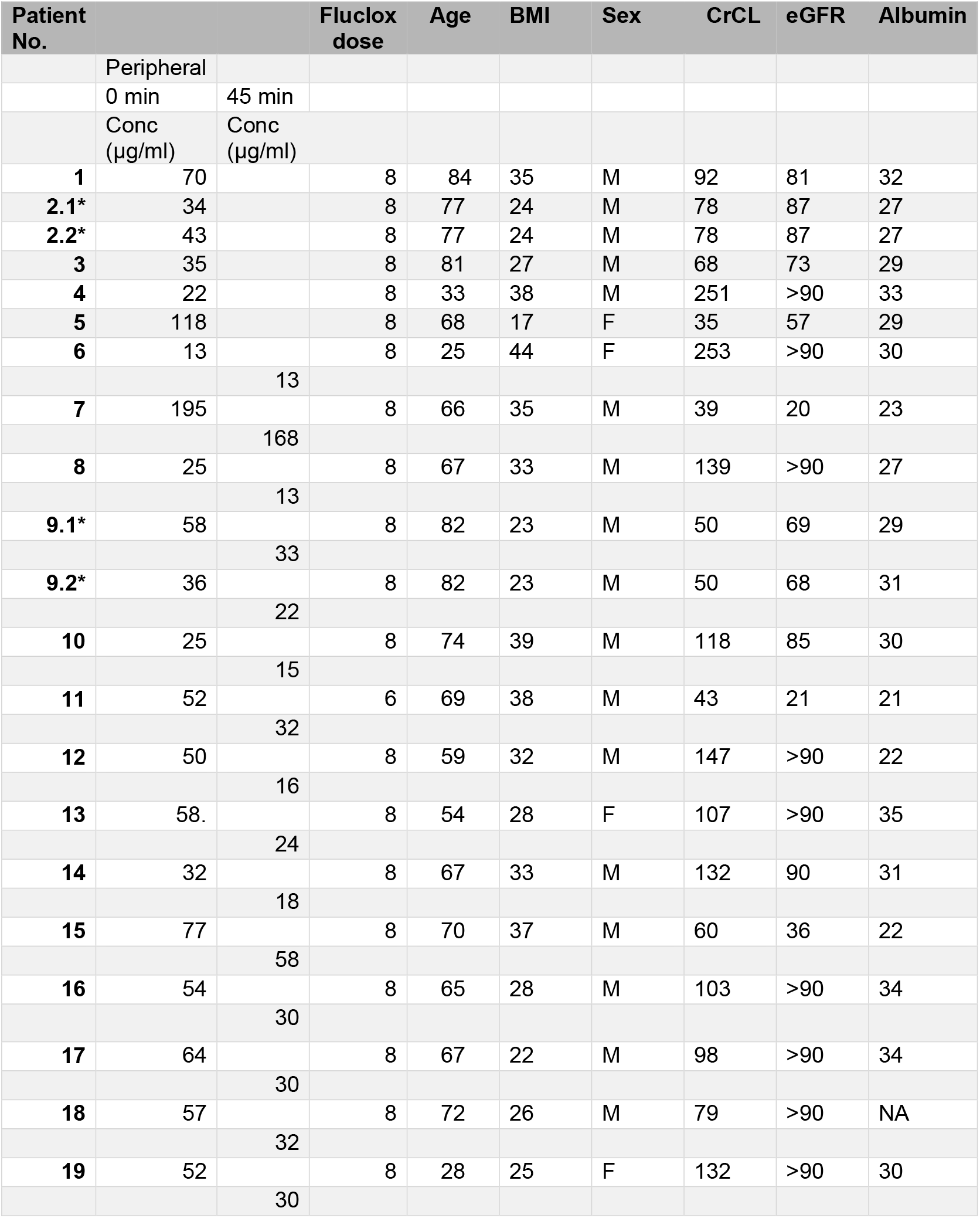
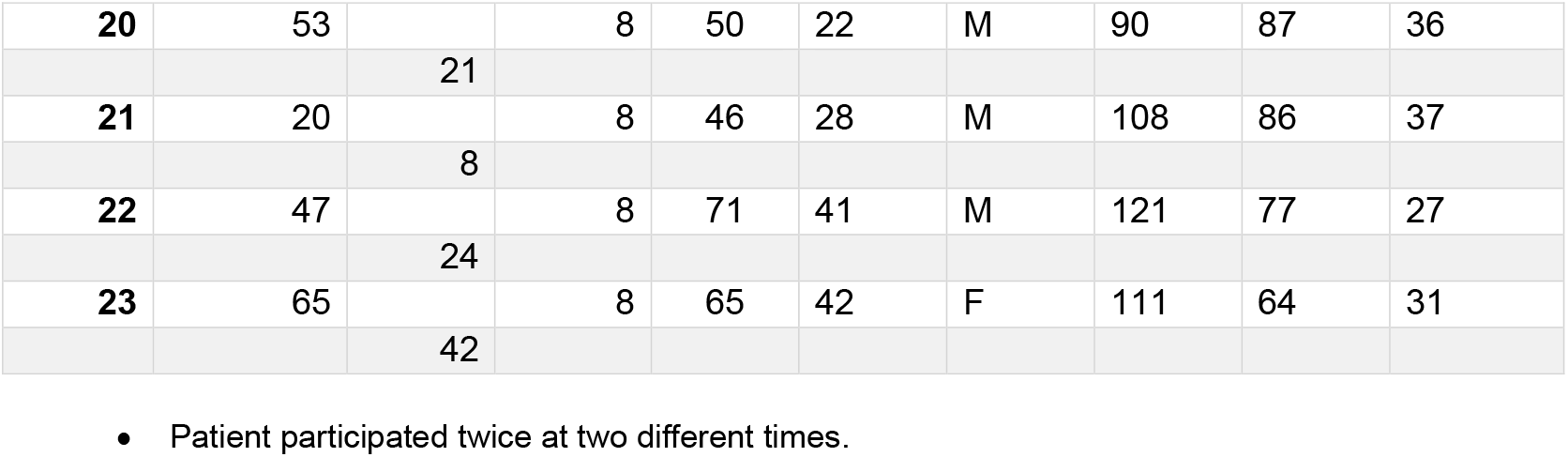
Basic characteristics and drug concentrations of 23 participants

## Abbreviations

CrCL: Creatinine Clearance calculated with Cockcroft-Gault equation
GFR: Glomerular filtration rate
OFV: Objective function value

